# Residue-dependent transition temperatures and denaturant midpoints in the folding of a multi-domain protein

**DOI:** 10.1101/2022.10.08.511446

**Authors:** Zhenxing Liu, D. Thirumalai

## Abstract

As a consequence of the finite size of globular proteins, it is expected that there should be dispersions in the global melting temperature (*T*_*m*_) and the denaturation midpoint (*C*_*m*_). Thermodynamic considerations dictate that the dispersions, Δ*T*_*m*_ in *T*_*m*_ and Δ*C*_*m*_ in *C*_*m*_, should decrease with *N*, the number of residues in the protein. We performed coarse-grained simulations of the Self-Organized Polymer (SOP) model of the multi-domain protein, Adenylate Kinase (ADK) with *N* = 214, in order to calculate thermal and denaturation unfolding titration curves. The results show that 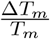 and 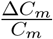 are non-zero and follow the previously established (*Phys. Rev. Lett*. **93** 268107 (2004)) thermodynamic 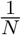 scaling for proteins accurately. For ADK, the dispersions are small (≈ 0.004), which implies that the melting temperature is more or less unique, which is unlike in BBL (*N* =40) where 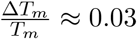.

## Introduction

In a largely unnoticed paper in 1997 Holtzer *et al*.,^1^ used ^13^*C*_*α*_ chemical shifts to show that the melting temperature depends on the residue number. In particular, the melting temperatures at 12 labelled sites in a 33-residue GCN4-like zipper varied in the range ≈ (42°*C* − 50°*C*). About five years later, we rationalized^2^ the Holtzer findings using lattice model simulations, thermodynamic arguments based on finite size scaling, and linked the variations in residue-dependent melting temperatures to local cooperativity. In other words, both sequence and local environment determine the extent of melting temperature variations with respect to the mean. In a most insightful study, Sadqi, Fushman and Munoz used proton NMR experiments on the forty-residue protein BBL^3^ and measured 158 local proton melting curves (backbone, side chain, and amide). They found that the distribution of the melting curves is roughly a Gaussian with a mean (*T*_*m*_) of 308K and dispersion, Δ*T*_*m*_ ≈ 17K (Figure 2b in the BBL study^3^). A related computational study on protein L,^4^ which used denaturants as a perturbation, showed that the midpoint transition is residue-dependent. Collectively, these reports have established that neither the melting temperature nor the denaturant midpoint in the equilibrium folding of proteins is unique. Similar findings have also been reported for RNA in which the midpoint concentration of divalent ions depends on the location of the nucleotide.^5^ From a theoretical perspective the residue-dependent ordering is best explained as a consequence of finite size effects, although this aspect is not appreciated in the the protein folding community.

**Figure 1:**
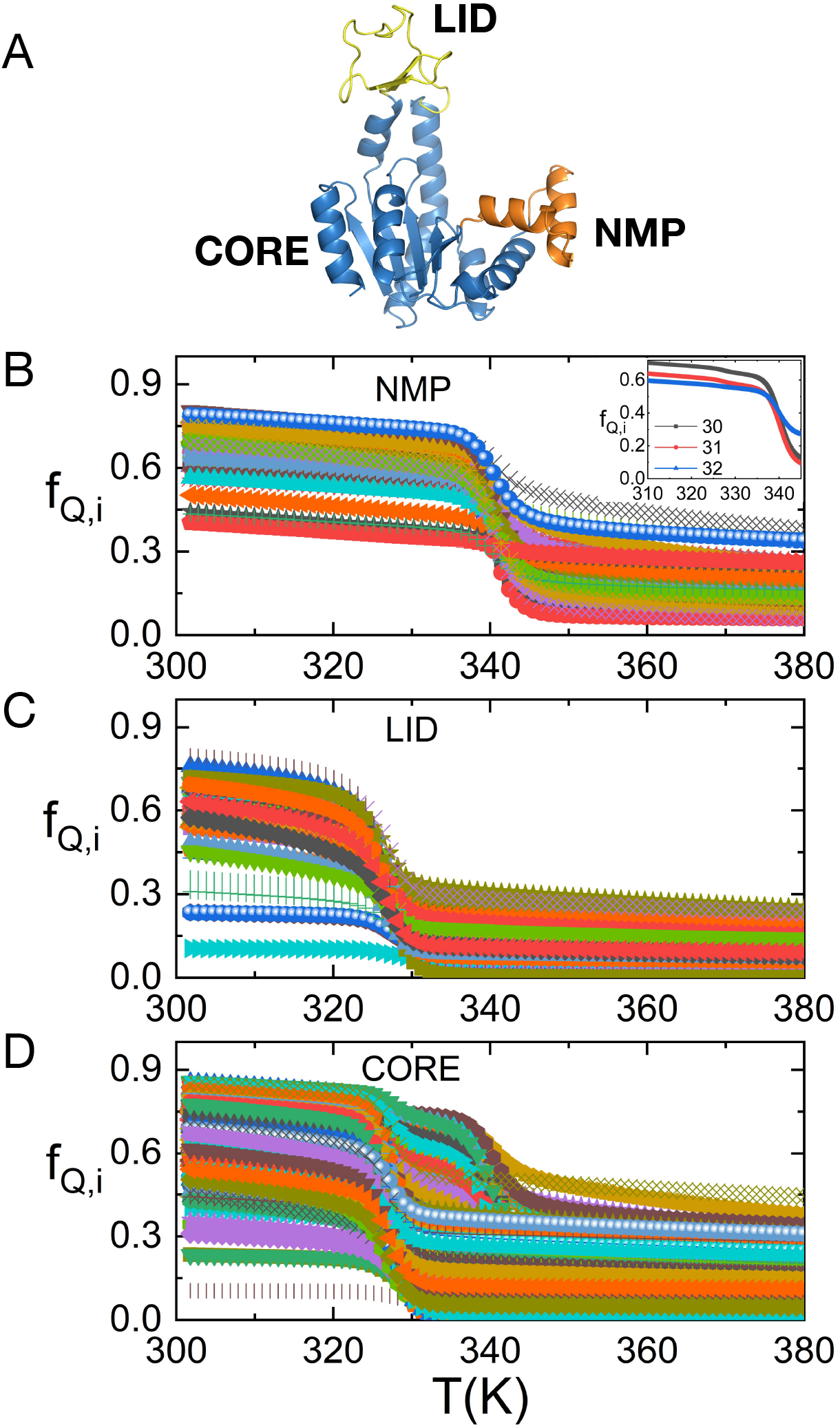
(A) Cartoon representation of ADK. The domains are also labeled. (B-D) Temperature-dependent fraction of native contacts formed for each residue in the NMP domain (B), the LID domain (C) and CORE domain (D). The inset in (B) show the dependence of *f*_*Q,i*_ on temperature for residues 30, 31, and 32 in the NMP domain.

**Figure 2:**
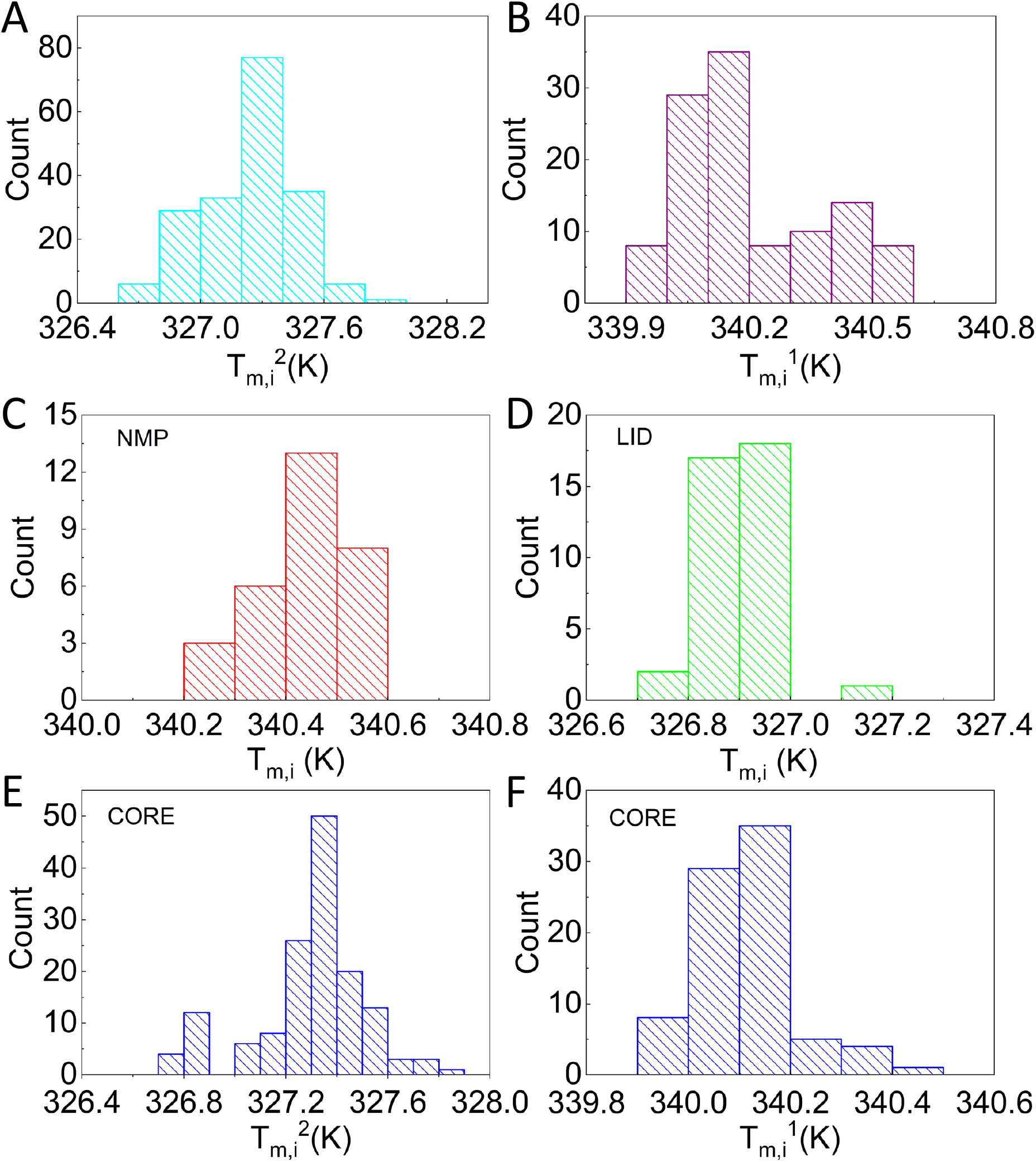
Distribution of transition temperatures around 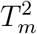 (A) and around 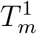 (B) for all 214 residues. Distribution of transition temperatures for residues in NMP domain (C), LID domain (D) and CORE domain (E,F).

That there ought to be variations in the local melting temperatures follows from standard thermodynamic arguments applied to finite sized systems. Several years ago, we showed, by analyzing experimental data for over 30 proteins (with *N* ≤ 150 where *N* is the number of amino acid residues) that 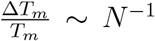 (Figure 3 in Ref.^2^ and Figure 2 in Ref. ^6^). For instance, using the results in Figure 2 in the previous report,^6^ we find that the expected value for 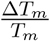 is ≈ 0.03 for BBL (*N* = 40) which is close to the measured value.^3^ The finite size analysis also predicts that as *N* increases Δ*T*_*m*_ should decrease.

**Figure 3:**
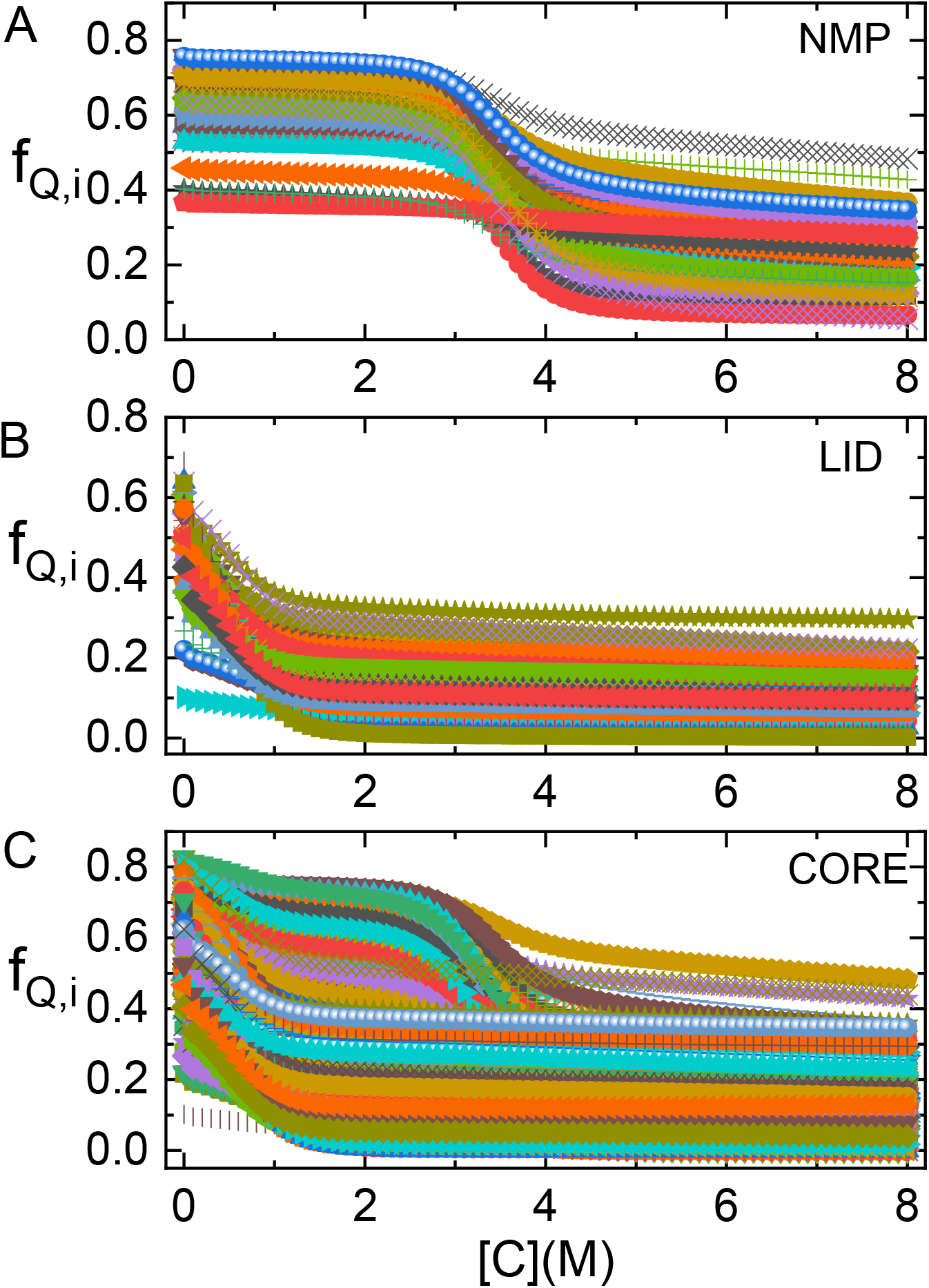
Residue-dependent ordering as a function of [GdmCl], measured using the fraction of native contacts formed for each residue. (A) NMP domain, (B) LID domain, and (C) CORE domain.

In this note, using of the Self-Organized Polymer (SOP) model^7,8^ we calculated both the global and residue-dependent folding of Adenylate Kinase (ADK) as a function of temperature and concentration of guanidinium chloride (GdmCl). Folding of ADK, which is a phoshotransferase enzyme with three domains (NMP, LID, and CORE), has been extensively investigated using experiments and simulations. ^9–13^ We find that the dispersions in the melting temperature 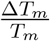 and the corresponding denaturant-induced dispersions in the midpoint 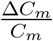 (*C*_*m*_ is the global midpoint of unfolding) are small because *N* = 214 for ADK is big enough for finite size effects to be negligible. An implication of this finding is that when *N* exceeds about 200 finite size corrections to the global unfolding temperature and the transition midpoint are minimal.

## Methods

### Coarse-Grained Model

We used the SOP-SC (Self-Organized Polymer-Side Chain) model for the protein.^7,8^ Each amino acid residue is coarse grained into two interaction centers with one at the C_*α*_ position and the other at the center of mass of the side chain. The SOP-SC energy function is

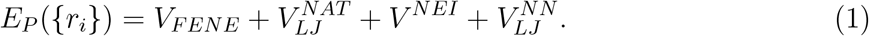

The detailed functional forms for 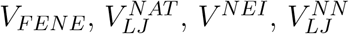 are described elsewhere.^8^

### Molecular Transfer Model

We used the Molecular Transfer Model (MTM) to take the effect of denaturants into account. In the MTM, whose theoretical basis is provided in a previous studies,^14,15^ the effective free energy function for a protein in aqueous denaturant solution with concentration [*C*] is given by,

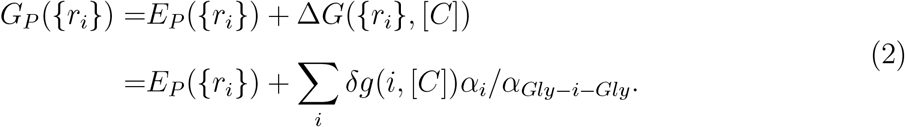

In Eq.(2), Δ*G*({*r*_*i*_}, [*C*]) is the free energy of transferring a given protein conformation from water to an aqueous denaturant solution, *δg*(*i*, [*C*]) is the transfer free energy of the interaction center *i, α*_*i*_ is the solvent accessible surface area (SASA) of the interaction center *i*, and *α*_*Gly−i−Gly*_ is the SASA of the interaction center *i* in the tripeptide *Gly* − *i* − *Gly*. We used the procedure described elsewhere^16^ to calculate the thermodynamic properties in the presence of denaturants.

### Langevin Simulations

Assuming that the dynamics of the protein is governed by the Langevin equation, we performed simulations using a low friction coefficient. ^17^ The equations of motions were integrated using the Verlet leap-frog algorithm. We exploited Replica-Exchange Molecular Dynamics (REMD)^18–20^ to enhance conformational sampling.

## Results

### Residue-dependent melting temperature,T_*m,i*_

The temperature dependence of the fraction of native contacts formed by the *i*-*th* residue, f_*Q,i*_, shows that each residue has one or two transition temperatures (Figure 1), depending on the domain. The transition temperature for each residue, T_*m,i*_, is identified by the peak position in 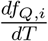. From the the distributions of 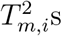 and 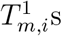 in Figure 2A and B, we find that Δ*T* for 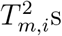 and 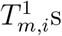 are 1.4K 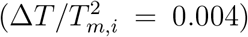 and 0.7K 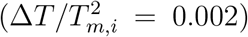, respectively. Thus, thermal dispersion in the melting temperature is small. Analysis of the data for 32 proteins^6^ showed that 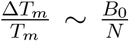 where *B*_0_ is close to unity. For *N* =214, the scaling relation predicts that 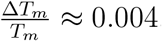, which accords well with the simulation result.

Our previous simulations showed that there are two global melting temperatures (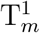 and 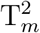 with 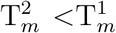) for ADK. For the NMP domain, only residues 30,31,32 have two distinct melting temperatures. The lower melting temperatures for these tree residues are T_*m,i*_=327.4K, 327,4K, 327.3K, respectively, which are around the global 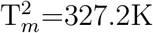 that was reported in an earlier study.^8^ The higher melting temperatures for all the residues are found to be T_*m,i*_=340.2K, which are nearly identical to the global 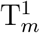. All other residues in NMP domain have a unique melting temperature, T_*m,i*_, close to the global 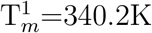. The distribution of the T_*m,i*_ around 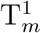 shows that the T_*m,i*_s are a little above 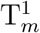 (Figure 2C). For the LID domain, all the residues melt at T_*m,i*_ that is close to the global 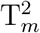. The T_*m,i*_ distribution (Figure 2D) shows that they are a bit below the global 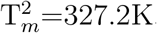. For the CORE domain, some residues have one melting temperature whereas others have two.The values of T_*m,i*_s cluster around the global 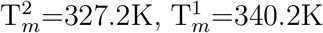, respectively (Figure 2E and F).

### Residue-dependent denaturation midpoint C_*m,i*_

The denaturant-dependent fraction of native contacts for the *i*-*th* residue, f_*Q,i*_, shows that each residue has one or two midpoint concentrations (Figure 3). We associate the midpoint concentration, C_*m,i*_, for the *i*-*th* residue with the peak position in 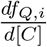. The distributions of 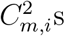 and 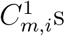 are shown in Figure 4A and B. The values of Δ*C*_*m*_ for 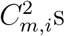 and 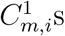 are 0.14M 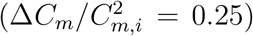 and 0.16M 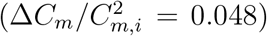, respectively. Interestingly, there are larger variations in the dispersions when unfolding is induced by denaturant compared to thermal unfolding. Analysis of experimental data led to a similar conclusion in a previous study. ^6^

**Figure 4:**
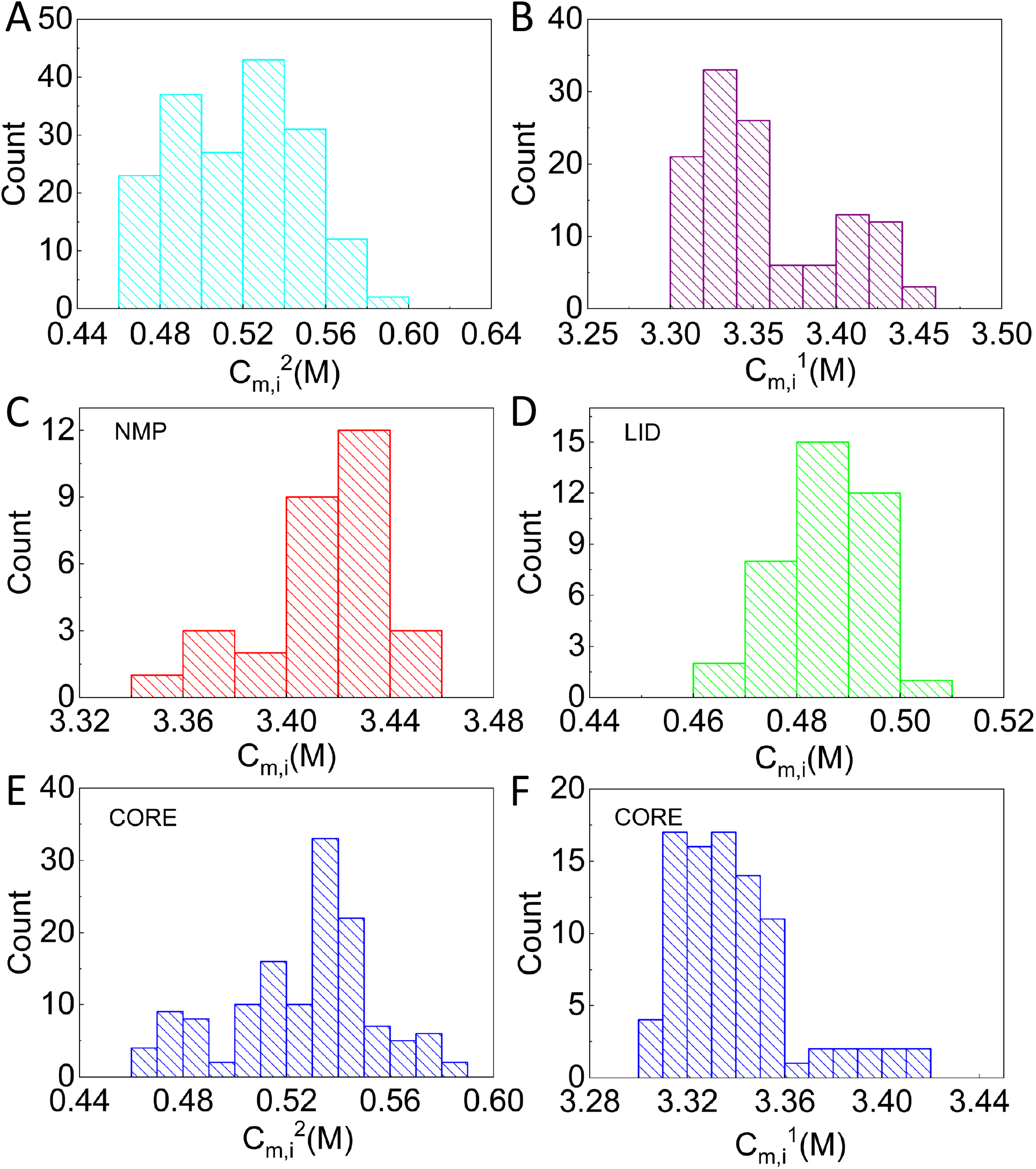
Distribution of midpoint concentrations around 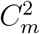 (A) and around 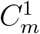 (B) for all 214 residues. Distribution of midpoint concentrations for residues in NMP domain(C), LID domain (D) and CORE domain(E,F).

**Figure 5:**
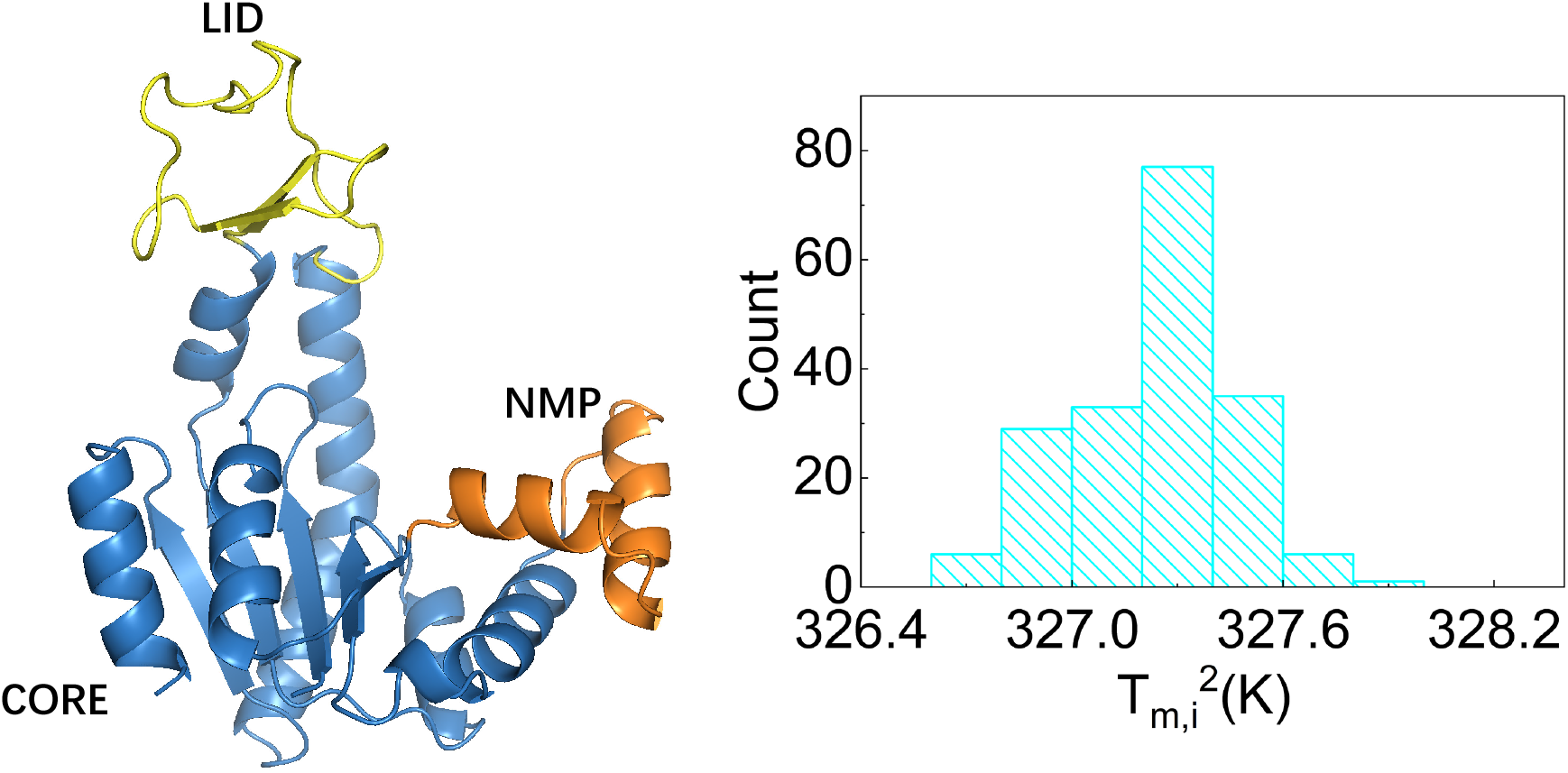
Table of Contents

Just as in the case of thermal folding, in the NMP domain, all the residues (except residues 30,31,32 that have two transition midpoints with the lower one at C_*m,i*_=0.55M, 0.56M, 0.54M, respectively. These values are close to the global midpoint concentration, 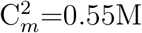.) unfold at 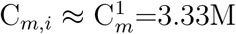. The distribution of the C_*m,i*_ shows that the C_*m,i*_s are a little above 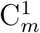 (Figure 4C). In the LID domain, all the residues unfold at C_*m,i*_ around the global 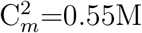. The distribution in Figure 4D shows that they are moderately below the global 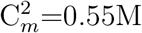. For the CORE domain, some residues have one C_*m,i*_ and others have two C_*m,i*_s. The distribution of their C_*m,i*_s (Figure 4E and F) cluster around the global 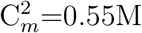 and 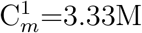, respectively. Thus, qualitatively the variations in the denaturant midpoints and thermal melting temperatures exhibit similar behavior.

## Conclusions

The present study, which extends our previous works^2,6,21^ that investigated finite size effects on folding of globular proteins, shows that as the length of the protein increases the dispersions in the melting temperature or the denaturation midpoint decrease in accord with finite size scaling.^6^ The dispersions in the melting temperature follow the predicted scaling, 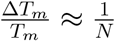. Simulations on ADK, a larger protein, confirm the theoretical predictions, which is a general consequence of thermodynamics of small systems. Because the dispersions are related to the extent of cooperativity,^2^ we surmise that the universal scaling for a measure of cooperative transition (see Figure 3 in Ref. ^6^) should also hold. Finally, denaturation by pressure also results in residue-dependent unfolding. It has been shown^22^ that different residues unfold at different values of pressure (see Figure 6c in Ref.^22^). Unfolding by any form of perturbation (temperature, denaturant, pressure and mechanical force) should result in residue-dependent response, reflecting the nature of the local environment. It is possible to verify these predictions using precise single molecule experiments.

## Acknowledgements

This work was supported by the National Science Foundation (CHE 19-00033) and the Welch Foundation through the Collie-Welch Chair (F-0019). ZL acknowledges financial support from the National Natural Science Foundation of China (11104015, 11735005).

